# Extracellular space diffusion modelling identifies distinct functional advantages of archetypical glutamatergic and GABAergic synapse geometries

**DOI:** 10.1101/2025.10.29.685288

**Authors:** Paula Gimenez, Rohisha T. Shakya, Fidel Santamaria, Jan Tønnesen

## Abstract

The brain extracellular space (ECS) is a convoluted compartment of nano- and microscale interconnected ducts. A key step in signaling between neural cells is diffusion of signaling molecules through the ECS, yet, signaling is generally considered solely from the stance of cells and their properties. Where ECS diffusion is addressed, this is commonly done using volume-averaging techniques blind to individual signaling events and ECS geometry. We hypothesized that ECS geometry can shape local diffusion and thereby tune signaling arising from point-sources. To access the scale of individual transmitter release events and synapse geometries, we developed a computational diffusion model, *DifFlux*, based on super-resolved images of hippocampal ECS in live mouse brain slices and combined this with single molecule Monte Carlo diffusion simulations. Our approach allows us to simulate diffusion of molecules of our choosing in true live ECS geometries. We asked how the ECS shapes local diffusion in dense neuropil and along larger cellular processes in CA1 *stratum radiatum*. We observed local diffusional anisotropy and directionality imposed by ECS geometry. Further, we identified distinct functional advantages of dendritic spine and somatodendritic synapse ECS geometries, shedding light on the longstanding conundrum of why glutamatergic and GABAergic synapses are so conspicuously morphologically different. Our modelling broadly identifies ECS structure as a direct modulator of extrasynaptic signaling that can operate in parallel to conventional regulation mechanisms.

## Introduction

The brain extracellular space (ECS) is a geometrically complex reticular continuum filled with interstitial fluid and stationary extracellular matrix (ECM) components (Kamali-Zare and Nicholson, 2013). It mediates diffusional transport of signaling molecules, ions, and metabolites to and from cells (Syková, 2004). Yet, despite its fundamental importance, it remains unknown if the ECS geometric structure modulates diffusion across any scale in a manner that facilitates specific physiological functions (Özçete et al., 2024). This is because existing knowledge has been provided mostly through experiments on the scales of seconds and hundreds of microns [For reviews, e.g. (Syková and Nicholson, 2008; Tønnesen et al., 2023)], though this is far from the range of individual point-source transmitter release events that are the basis of neuronal signalling and which unfold on millisecond and nano- to micrometer scales (Mauk and Buonomano, 2004). Diffusion experiments on coarser scales inevitably volume-average data across space and time, and any information about biological variation around the average remains inaccessible. There is a particular lack of knowledge on how neurotransmitters may undergo local anisotropic diffusion in the ECS before finding their target, and accordingly whether ECS geometry serves a role in signal modulation. This question becomes even more pertinent considering the diffusional properties of the ECS change with aging, trauma, and pathology (Syková and Nicholson, 2008), which are conditions associated with changes in signaling and neuronal plasticity in general.

The two archetypical ionotropic neurotransmitters of the mammalian brain are excitatory glutamate and inhibitory gamma-aminobutyric acid (GABA). Glutamatergic signalling through independently tunable AMPA receptor rich glutamatergic synapses operates with typical deactivation time constants around three milliseconds and is considered at the core of neuronal wiring of cognition and learning (Kennedy, 2016). Outside the synapse glutamate still serves as a transmitter, though extra-synaptic glutamatergic receptors serve a less clear-cut role and is not found to be a strong general modulator of excitability (Pál, 2018). GABA-A receptors activate synaptic conductances with deactivation time constants of tens of milliseconds to cause fast, phasic inhibition (Farrant and Nusser, 2005). Additionally, GABA spills over from the synapse and diffuses to peri- and extra-synaptic areas where it gives rise to a tonic inhibitory current mediated also primarily by GABA-A receptor subtypes (Agnati et al., 1986; Rusakov and Kullmann, 1998; Farrant and Nusser, 2005). Extrasynaptic tonic GABAergic inhibition operate across cell types and brain regions where it amounts to 30-50% of total inhibition and ubiquitously sets baseline excitability (Brickley et al., 1996; Stell and Mody, 2002; Cope et al., 2005). This identifies the two main synaptic transmitter systems as not merely functionally opposite, but also differently implemented and regulated in terms of time scales and extra-synaptic effects.

In addition to the functional differences, the structural geometries of GABA and glutamatergic synapses differ conspicuously. While the pre-synaptic sides appear largely similar as axonal boutons, the post-synaptic sides differ and the differences are conserved across species and cell types, which suggests a physiological advantage (Gray, 1959; Santuy et al., 2018). On most neurons, including hippocampal CA1 pyramidal cells, glutamatergic post-synapses are located on dendritic spines protruding to offset the synapse from the surrounding dendritic membrane. By contrast, GABAergic post-synapses predominantly form directly on the soma or dendritic trunk and are level with the surrounding membrane (Megías et al., 2001; Kwon et al., 2019). Dendritic spines compartmentalize signaling inside the spine head, which adds to their independent functional plasticity and to the computational power of the brain (Sala and Segal, 2014). Conversely, it remains an enigma whether there is a biological advantage of membrane level somatodendritic GABAergic post-synapses and why these two transmitter systems are so consistently morphologically different.

We hypothesized that the respective geometries of excitatory and inhibitory synapses will differently modulate perisynaptic diffusion and spill-over to neighboring synapses and extrasynaptic receptors. Specifically, we propose that diffusional clearance from spines is facilitated so that synaptic crosstalk is minimized, whereas somatodendritic synapse geometry by design enhance lateral diffusion to nearby extrasynaptic receptors to enhance tonic inhibition. To test this hypothesis, we used fluorescence super-resolution shadow imaging (SUSHI) to image the ECS and neuropil geometry in live mouse brain slices (Tønnesen et al., 2018). We first analyzed the ECS volume fraction, α, and spatial frequencies, and then built a computational diffusion model to simulate diffusion in SUSHI images and test how these parameters could impact perisynaptic diffusion. The model hinges on Fick’s law stating that diffusional fluxes through a compartment scale with its cross-sectional area. We extrapolate this to SUSHI image pixels and determine to what extent these are a diffusionally accessible part of the ECS or whether they are instead occupied by cellular structure. This allows analyses of diffusion from simulated glutamatergic and GABAergic synapses into their perisynaptic ECS. Lastly, we simulate increasing frequencies of synaptic GABA activity through somatodendritic synapses to learn if regular point-source release could result in steady state micro-gradients of GABA along the cell membrane and facilitation of local extrasynaptic tonic inhibition.

The combination of super-resolution imaging in live brain slices and computational diffusion modelling in the subsequent images provides a unique opportunity to investigate perisynaptic diffusion following individual release events at single at nano- and micrometer scales, beyond the reach of volume-averaging approaches. Our results points to the ECS as a signal regulator with transmitter system specific functional roles for respective glutamatergic and GABAergic synapses.

## Methods

### Organotypic mouse brain slices

We prepared organotypic hippocampal slices from postnatal day 5-7 C57BL/6J mice, as previously reported in detail (Tønnesen et al., 2018) (<u>protocols.io</u>). In brief, hippocampi were sliced at 250 μm on a vibratome and embedded in a plasma-thrombin clot on a 12 x 24 mm glass coverslip (Thickness #1) before culturing in a roller-drum incubator at 10 RPM in tubes with 0.75 ml medium consisting of 50% MEM, 25% horse serum and 25% HBSS supplemented with 12 mM glucose and penicillin/streptomycin (Gähwiler, 1988). Cultures were imaged after 2-4 weeks of culturing. All experimental procedures were performed in adherence to Spanish law (Royal decree 53/2013, BOE 08-02-2013) and European Community Council Directive (2010/63/EU). Experiments were prospectively approved by the ethics committee of the University of the Basque Country (M20/2021/339).

### Super-resolution STED and SUSHI imaging in live brain tissue

We used a custom-built microscope based on an inverted body (DMi8, Leica) with an 93x and 1.3 numerical aperture (NA) glycerol immersion objective with a motorized aberration correction collar (Leica). The STED beam was derived from a 592 nm 500 ps pulsed laser (Katana 06, NKT Photonics), and the excitation beam from a 485 nm 90 ps pulsed laser (Quixx, Omikron). These were brought into temporal phase overlay using the built-in pulse- sync and pulse-delay module of the 485 nm laser. Resolution enhancement was achieved by splitting the STED beam and sending the two parts through a 2π vortex phase plate (doughnut PSF) and a 1π helical phase plate (bottle beam PSF; both plates from Vortex Photonics) before recombining them to achieve x,y,z resolution enhancement of the emission PSF (Tønnesen et al., 2018). We primarily imaged with majority power in the bottle beam PSF for z-resolution enhancement and using STED power of around 15 mW in the objective back aperture, adjusting the settings to the given imaging session. The 15 mW STED power typically provide a volume resolution around 125 x 125 x 300 nm^3^ on our setup, and we would tune power up or down during individual imaging sessions. The laser beams were scanned in x,y by means of a Yanus IV galvanometric scan head (ThermoFisher), and the back-scanned fluorescence was detected using an avalanche photo-diode detector (APD, Laser Photonics) coupled to a 50 µm multimode fiber acting as an optical pinhole for confocal detection (Thorlabs). Z-axis scanning was performed using a piezo-focuser (PIFOC, Physik Instrumente). The microscope was controlled through an FPGA-based data acquisition card using designated Imspector software (Abberior Instruments).

For imaging on our inverted STED microscope, a glass coverslip with a slice culture was glued onto a custom-made polycarbonate imaging chamber using UV glue, so that the glass coverslip effectively became the chamber bottom to allow single-interface imaging from beneath. The slice imaging chamber was perfused with modified artificial cerebrospinal fluid (maCSF) containing sucrose 195 mM, KCl 2.5 mM, NaH2PO4 1.25 mM, NaHCO3 28 mM, CaCl2 0.5 mM, L-ascorbic acid 1 mM, pyruvic acid 3 mM, glucose 7 mM, and MgCl2 7 mM, at 32-35°C. To visualize the ECS we used the protocol described in (Tønnesen et al., 2018). In brief, we added 20-30 µM calcein to the perfusion solution, which distributes homogeneously in the ECS without entering cells and allows super-resolution STED imaging of the neuropil. For 2-color SUSHI images of dendritic spines in their neuropil context we used previously published images from (Tønnesen et al., 2018) with permission. Images were minimally processed by applying a 1-pixel medial filtering to remove single pixel noise arising from photo-detector dark counts (**Figure 1A**). The image fluorescence signal histogram was normalized by adjusting for background, *Fbckg*, by imaging inside somas devoid of fluorophores, and for the maximum observed value, 𝐹_𝑚𝑎𝑥_, obtained by imaging a fluorophore filled ECS volume. Each pixel in a given intensity normalized 8-bit image would accordingly have a value

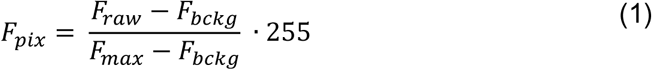

**Figure 1.**
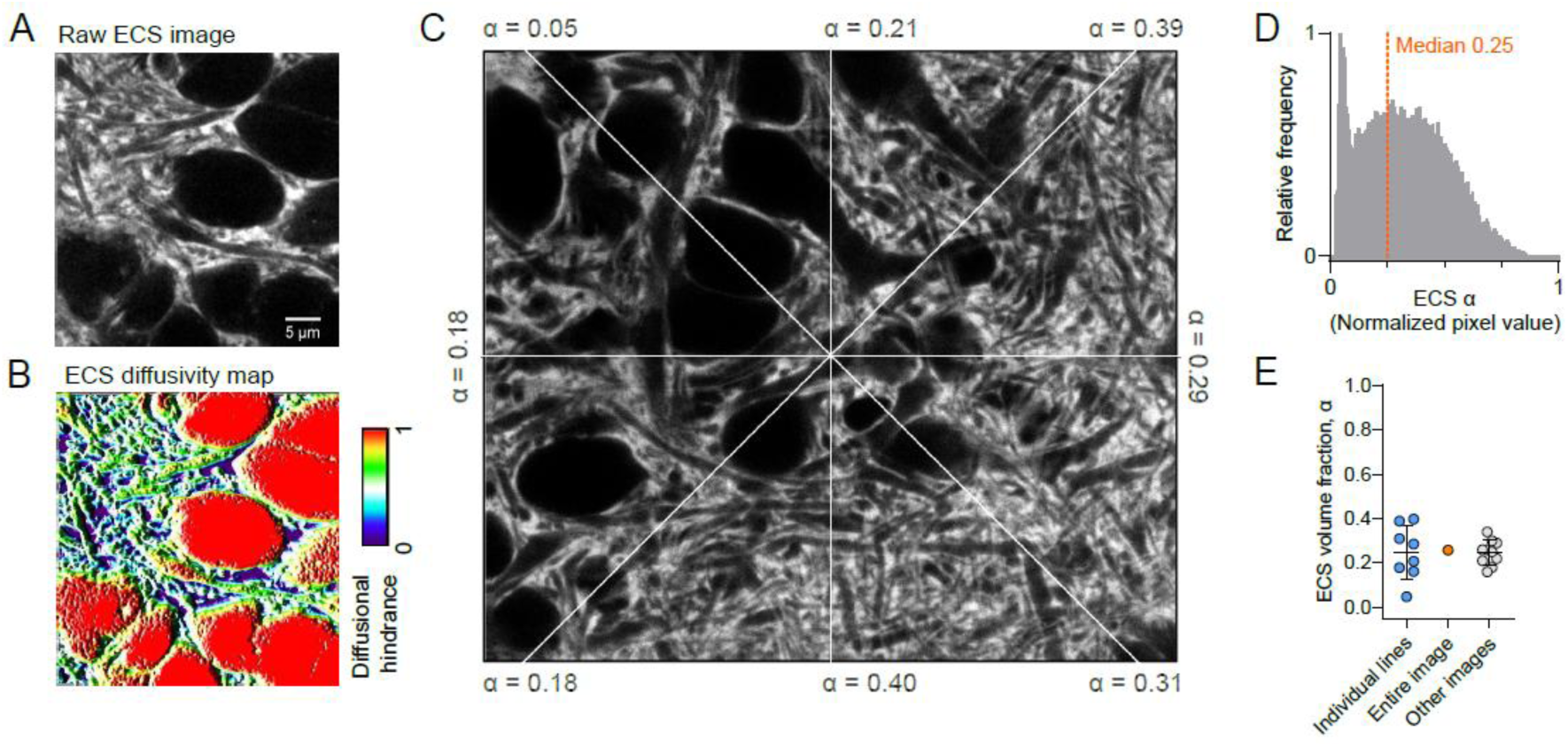
Diffusion model and local geometric and diffusional properties of ECS (**A**) Our diffusion model utilizes the ECS structural information of SUSHI images to effectively build a diffusional accessibility map (**B**). (**C**) Line profile analysis of radial lines from the image center returns widely varying α values between 0.05 and 0.4. (**D**) Intensity-normalized pixel histogram of image in (C), where pixel values directly correspond to α. The median of 0.25 is depicted by dotted line. (**E**) Calculated α for line profiles and image in (C) as well as for other images (n = 10 images, 8 slices, 6 animals; Mean and SD).

Where *Fraw* is the unadjusted raw pixel value of that pixel.

### Modelling ECS diffusion fluxes on nanometer and microsecond scales

Diffusional fluxes in a homogeneous medium scale with the diffusion coefficient, *D*, of the observed particle which is spatially confined by the delineating compartment borders. To model ECS diffusion we assumed an ECS filled with homogeneous interstitial fluid resembling saline and delimited structurally by the ECS geometry. In this scenario diffusion is free in pixels filled with pure 100% ECS, whereas diffusion cannot take place in pixels representing pure cellular structure (0% ECS). Diffusion is partially obstructed in pixels partly occupied by cellular structure (0% < ECS < 100%) (**Figure 1B**). The ECS geometry- weighted effective diffusion coefficient in a given pixel, 𝐷_𝑝𝑖𝑥_, will thus be a function of the pixel ECS volume fraction, 𝛼_𝑝𝑖𝑥_ that can be easily read out as the relative pixel intensity *Fpix*

We assume a linear mapping from the calculation 𝛼 to 𝐷_𝑝𝑖𝑥_:

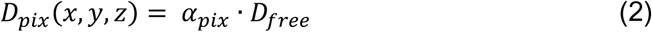

Our model was built in MATLAB (Natick, MA) based on a diffusion equation for mass transport in porous brain tissue (Nicholson and Phillips, 1981; Nicholson, 2001) assuming no advective interstitial fluid bulk flow.

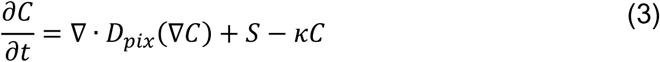

Where 𝐶 is the concentration, 𝑆 is the source magnitude, defined as the amount of substance delivered per unit time and 𝜅 is the non-specific clearance coefficient (Arranz, 2014). We used this coefficient to consider diffusion of molecules in the z-axis (kz) and thus build a simplified 3D model to model diffusion in 2D images.

In point source release simulations 𝑆 is the number of particles per volume unit at time zero:

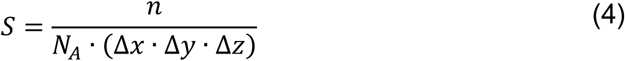

Where *n* is the number of particles, *NA* is Avogadro’s number and Δx, Δy, Δz are the image pixel or voxel sizes. For all our simulations we assumed leaky boundary conditions defined by an escape factor (*ef*) between 0.7 and 0.98 chosen to mimic a surrounding infinite field of ECS. We solved the diffusion equation numerically using the Forward Euler Method and with time steps on the scale of 1 µs, adjusted for individual images.

Simulations were performed in 2D and 3D as indicated. Further, to reduce the computational cost of 3D simulations and increase the applicability of our model we developed a simplified 3D diffusion model approach, *3Dsmpl* to model diffusion in x,y image frame without specific knowledge about the above and below image z-planes and still allowing escape from the plane. To account for 3D diffusional escape out of a given SUSHI x,y plane we introduce a clearance coefficient, 𝜅 = α ∗ 𝜅(𝑡), so that clearance is higher at pixels with higher volume fraction.

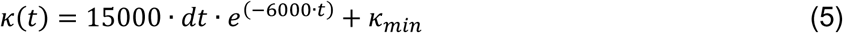

Where, 𝜅_𝑚𝑖𝑛_ = 75 · 10^6^ · 𝑑𝑡^2^ was observed to work well across our images (**Suppl. Fig. 1**). For simplicity we refer to our extracellular space diffusion flux model as *DifFlux*.

### Monte Carlo diffusion modeling in live ECS geometries

The diffusion of single particles is defined by their probability, 𝑝, to randomly displace from one location to another at every Δ𝑡 . Like in the *DifFlux* approach, for Monte Carlo simulations we assumed a linear mapping of 𝑝 as a function of 𝛼(𝑥, 𝑦, 𝑧) in a given SUSHI image, again exploiting the ECS structural information available in each pixel. At every step a random number is used to select an axis of movement with equal probability, 𝑝_𝑑𝑖𝑟_ = ^1^ 𝑜𝑟 ^1^, depending on the number of dimensions, 2D or 3D. Another random number chooses the direction along the axis with probability 𝑝_𝑖_=^1^. We then use a homogeneously distributed random number, 𝑟, to simulate bouncing from an obstacle in the new selected location, 𝑝_𝑏_ = (1 − 𝛼(𝑥, 𝑦, 𝑧)). If, 𝑟 < 𝑝_𝑏_then the particle stays in the original voxel, as described in our previous work (Santamaria et al., 2010). We assumed toroidal boundary conditions and non-interacting particles.

### Analysis and statistics

All modelling and analyses were performed in raw images subjected to an evenly applied 1-pixel median filter to remove single pixel detector noise. Figure images are not further processed. The ECS *α* for an image frame or line profile was calculated as the median pixel value divided by 255 (the value 255 corresponding to *α* = 1) in the intensity-normalized images. We tested data for normality and applied parametric or non-parametric testing and descriptive statistics as specified for results throughout. We considered probabilities less than 0.05 significant.

## Results

### The ECS volume fraction is highly heterogenous and imparts anomalous diffusion

The normalized fluorescence intensity value of each pixel in each image corresponds to the ECS volume fraction *α*. We found that *α* varied considerably within images as well as between images and brain slices. Radial lines from the center of the SUSHI image depicted in **(Figure 1C)** reported ECS α values varying 8-fold between 5% and 40% with a mean and SD of 0.25 ± 0.12 (n = 8 lines), in excellent agreement with the α = 0.26 based on integrating all pixels of the image **(Figure 1D)**. The mean α value again fits well with the average across other *stratum radiatum* images from different slices where *α* = 0.25 ± 0.06 (n = 10 slices, 6 animals) **(Figure 1E)**. The observation of widely different α values of line profiles emerging from a common center within a given image suggests that diffusion will also depend on the considered diffusion direction from a source point, so that a given tissue tortuosity is valid only for a given diffusion direction. While α varied widely within and between tissue areas, the average was robust around 25%, in agreement with literature values (Van Harreveld et al., 1965; Lehmenkühler et al., 1993; Korogod et al., 2015).

We further quantified the structural heterogeneity of the ECS by 2D radial Fourier analysis in live tissue SUSHI z-stacks (n = 113 planes, 7 image stacks/slices/animals) (Ruzanski, 2025). The radially averaged power spectral density (PSD) shows a smooth transition from low to high spatial frequencies indicating a continuum of ECS geometries, again corroborating earlier results of lognormal distributions (Godin et al., 2017; Tønnesen et al., 2018) (**Figure 2A**). Our analysis focused on spatial frequencies ranging from 0.13 µ𝑚^−1^, corresponding to large-scale features, to 2 µ𝑚^−1^, which reflect image noise as determined by analyzing dark regions in the images. Given the high spectral similarity within each stack, we averaged PSDs per stack and then computed a grand average across all stacks. We identified two regions in the averaged PSD that can be approximated by a power law, 𝑃𝑆𝐷∼1/𝑓^𝛽^ . The lower spatial frequencies, 0.13 − 0.5 µ𝑚^−1^, dominated by neuronal somas and large dendrites, with a fitted exponent of 𝛽 = 2.04 (95% 𝐶𝐼: 2.02 − 2.06). This corresponds to Brownian noise, indicative of large-scale, smooth, and correlated ECS structure similar to natural images (Field, 1987; Ruderman and Bialek, 1994). Such structure suggests long-range correlations akin fractals and disordered media (Havlin and Ben-Avraham, 1987; Hambly and Jones, 2000). The high frequencies (0.5 − 2 µ𝑚^−1^) were dominated by finer ECS structures, with a fitted exponent of 𝛽 = 3.62 (95% 𝐶𝐼: 3.58 − 3.66) . This value of 𝛽, sometimes called super-Brown or colored noise, reflects suppression of fine-scale detail, consistent with smoother extracellular boundaries or blurred fine structure, rather than sharp discontinuities. This may indicate a breakdown of fractal behavior at small scales potentially due to limitations in optical resolution which could smooth the images at small scales, masking actual fractal behavior. The PSD showed consistency between slices, suggesting a conserved ECS geometry within the CA1 region (**Figure 2A**). Our combined α and Fourier analyses show that the ECS geometry is highly variable yet conserved with comparable average α and geometry within the CA1 *stratum radiatum*.

**Figure 2.**
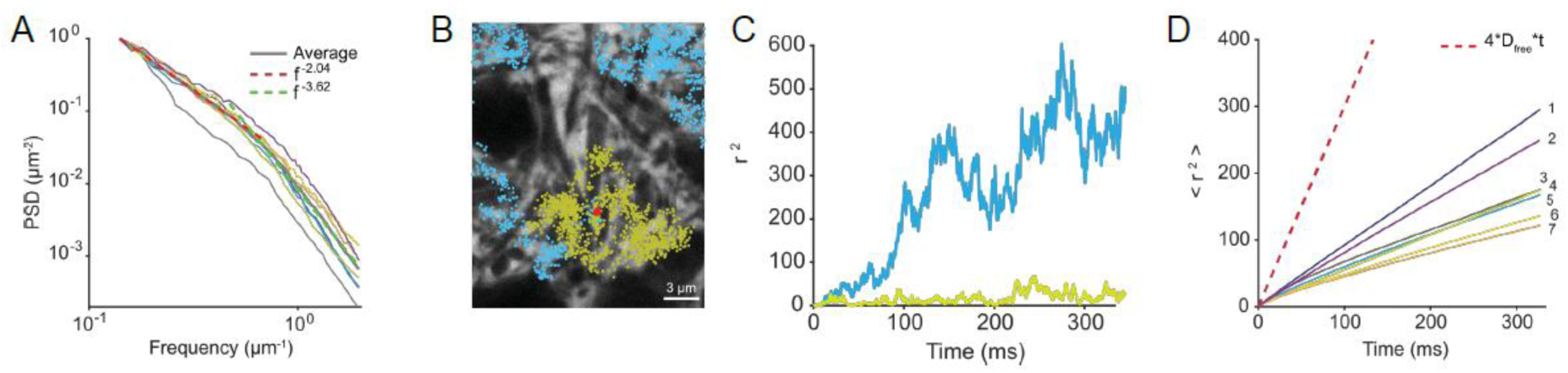
Monte Carlo simulations of ECS diffusion **(A)** Fourier based radially averaged power spectral density (PSD) analysis of ECS geometry from seven SUSHI image stacks shows a continuous distribution with apparent adherence to power law. The average PSD across all 7 stacks shown in black, the fitting exponent β = 2.04 (95% CI: 2.02- 2.06) (red dotted line) at low frequencies and β = 3.62 (95% CI: 3.58- 3.66) at high frequencies (green dotted line). **(B)** Two examples of Monte Carlo simulation of particle trajectories after release at a common source point (red), illustrating different resulting trajectories (blue and green) and **(C)** mean squared displacement (r^2^) values. **(D)** Average mean squared displacement (<r^2^>) values for middle plane of each of seven stacks stack illustrating anomalous diffusion relative to free diffusion (dotted line). A total of 50 random source point release events were simulated per image plane. A power law fit to the average <r^2^> plot yielded an overall exponent, γ, of 0.88 ± 0.02 SEM, n = 7.

To explore how ECS structural heterogeneity influences molecular diffusion we applied 2D Monte Carlo simulations of diffusion of a small molecule (Dfree = 0.5 μm^2^/ms). First, we observed that *DifFlux* and Monte Carlo simulations yielded comparable results when applied to simulate diffusion using similar parameters (**Suppl. Figure 1**). Using Monte Carlo simulations, we tracked the movement of 1,000 glutamate sized particles (*DGlu* = 0.75 μm^2^/ms)from 50 randomly selected source points over 300 ms using toroidal boundary conditions. Plotting the individual particle trajectories emerging from a given release site showed large variability in diffusion patterns, (**Figure 2B, C**). We calculated the mean squared displacement (MSD, < 𝑟^2^ >) for each source point to determine their average behavior. For each image frame, we averaged the MSD curves from 50 simulations. This showed slowed-down non-linear diffusion characteristic of anomalous diffusion (**Figure 2D**) (n = 7 frames/slices/animals). Indeed, a power law fit, <r^2^> ∼ t^γ^, where γ is the diffusion exponent, described the average MSD well with an overall value of 𝛾 of 0.88 ± 0.02 𝑆𝐸𝑀 (n=7 images). This deviation from normal diffusion (𝛾 = 1), was statistically significant in all cases (t-test and Wilcoxon, p<0.0001). Taken together, our Monte Carlo simulations corroborate that the *stratum radiatum* neuropil is geometrically heterogeneous, characterized by power-law geometric structure, and is associated with pronounced anomalous diffusion. Brain parenchyma anomalous sub-diffusion has been reported on volume-averaged scales though it has not been previously related to actual ECS geometries (Xiao et al., 2015).

### Tensor mapping identifies diffusion directionality based on ECS geometry

To further explore how ECS geometry could shape local molecular diffusion in the interstitial space we used *DifFlux* to simulate release of glutamate molecules (*DGlu* = 0.75 μm^2^/ms) from individual point sources in SUSHI images of dense CA1 neuropil (**Figure 3A**). We found that the normalized index of dispersion, corresponding to the variance, of resulting concentrations at given distances around the release site would be median 0.007 at 0.2 μm and reaching maximum of 0.047 at 2 μm then come down to become insignificant beyond three microns compared to all lesser distances (p < 0.001, n = 6 images, 4 slices/animals, Kruskal Wallis test; (**Figure 3B**). This shows that there is pronounced anisotropic diffusion at single micron scales immediately after release in *stratum radiatum* neuropil, though at larger scales the variation is canceled out. Functionally, this would mean that the ECS can have a stronger modulatory effect on diffusional processes on scales below a few microns.

**Figure 3.**
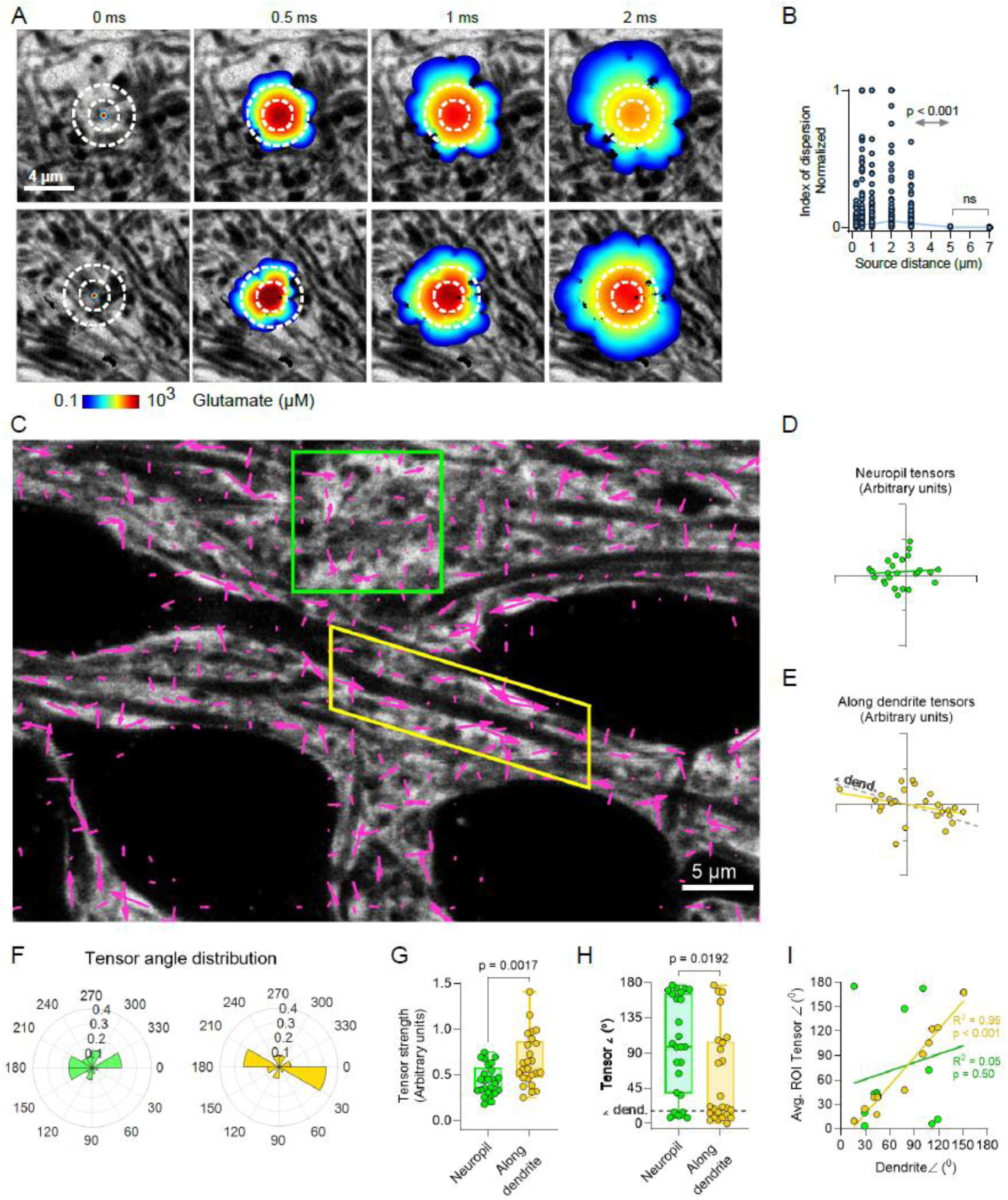
Local diffusion directionality is determined by ECS geometry (**A**) Two examples of DifFlux simulations of glutamate release in dense neuropil. Dotted circles shown for reference across time lapse. (**B**) Normalized index of dispersion of resulting glutamate concentrations at indicated source distances for the two examples in (A) and other slices illustrating that the variation is highest below 5 µm from the source point (n = 5 slices/animals, Kruskal-Wallis test). This confirms anisotropic diffusion at micron and submicron scales that diminishes at larger distances. (**C**) ECS diffusion tensor map showing local directionality (angle, and integrated tensor strength with goodness of fit (as length). (**D**) Tensors in dense neuropil (green box in C) appear scattered and random, (**E**) while along dendrites diffusion tensors appear to follow the direction of the extending dendrite. The dotted line indicates the dendrite extension angle. (**F**) Binned (30 degrees) directionality angle distribution of tensors show that in dense neuropil there is less clustering around specific angles than along dendrites where the vast majority of tensors fall in the bins0°-30° and 180°-210° that encompass the 16°/196° dendrite extension angle. (**G**) Comparing tensor strengths reveals that these are stronger along dendrites 0.66 ± 0.29 μm, mean ± SD) than in dense ECS neuropil (0.44 ± 0.16 μm, mean ± SD; Unpaired t-test, p = 0.0017). (**H**) Similarly, the 0-180° orientation of tensors significantly differed between dense ECS where tensors were distributed throughout the interval, and along dendrite where again the majority of tensors were clustered close to the 16° angle of the dendrite itself (dotted line). (**I**) analysis across experiments found that the angle of tensors along a dendrite would strongly correlate with the angle of the dendrite (R^2^ = 0.95, n = 12 frames, 10 slices, 6 animals), while in the same images tensors in dense neuropil would have a seemingly random distribution with respect to the dendrite (R^2^ = 0.05).

We next asked if point-source diffusion anisotropy was generally random or could translate into diffusional directionality across tissue areas, and modelled release of glutamate from 775 point-sources distributed in a regular grid pattern across an image. For each resulting diffusion point cloud we identified the furthest-from-source point and applied linear regression to identify this as main direction measured 100 µs after release (**Figure 3C**). The direction of each point was assigned a vector with a corresponding direction (angle) and magnitude (diffusion distance). We further incorporated the observed bias of the anisotropy by scaling the vector magnitude by a directionality coefficient to obtain a corresponding diffusion tensor. The directionality coefficient was determined by first calculating the distance to the source of all the points that describe the diffusion cloud perimeter for a given source point. Then we plotted the cumulative distribution of that distance for each cloud and estimated the area under the curve, with the normalized area under the curve corresponding to the directionality coefficient. Here, if the diffusion pattern goes toward isotropic with a circular spread the tensor magnitude goes toward zero and the tensor is short. Conversely, the stronger the anisotropy, the longer the tensor becomes.

Applying this strategy in a SUSHI image that includes both dendrites and dense neuropil, we found that ECS diffusion tensors appeared with low directionality in dense neuropil (**Figure 3D**), while along dendrite ECS diffusion tensors were stronger and following direction of the dendrite (**Figure 3E**). Accordingly, the majority of tensors were found in the direction bins 0°-30° and 180°-210° corresponding to the 16° (and accordingly 196°) angle of the dendrite in the field of view (**Figure 3F**). Comparing the respective neuropil and peri- dendritic tensor strengths (speed weighed by goodness of fit), neuropil tensor arbitrary values were 0.44 ± 1.6 (mean with SD, n = 25 tensors), while they were 0.66 ± 2.9 (n = 25 tensors) along the dendrite (*p* = 0.002, unpaired t-test) (**Figure 3G**). Tensor angles were also different between neuropil and dendritic ECS, with the neuropil median of 98 (interquartile range [IQR] 38-166) close to random (average 90°), whereas the dendritic ECS tensor median angle was 21° (IQR 9.4°-105°), close to the 16° angle of the dendrite (*p* = 0.192 Mann Whitney U test) (**Figure 3H**). This held true across experiments, where the dendritic angle would account for 95% of the variation in ECS tensor angles along dendrites and be significantly non-zero (Linear regression, *R^2^* = 0.95, slope 1.15, non-zero *p* < 0.0001, n = 12 frames, 10 slices, 6 animals), while in dense neuropil of the same images the corresponding *R^2^* was 0.05 and the 0.34 slope not different from zero *p* = 0.50 (**Figure 3I**). These results suggest that the structural geometry of the ECS shapes diffusion and that not only directionality but also diffusion speed is modulated, for example along dendrites.

### Glutamatergic synapse crosstalk from diffusional synaptic spillover

After observing local diffusional directionality imposed by ECS geometry, we tested if perisynaptic ECS geometry modulate extracellular synaptic crosstalk of glutamatergic synapses on spines. Single vesicular release of glutamate was simulated onto a post- synapse on a dendritic spine and the resulting concentrations at two neighboring spine synapses was analyzed (**Figure 4A**). Both neighbors were roughly 2 µm from the central source synapse and the spread of glutamate was simulated over 3 ms following release. We compared this SUSHI scenario to the same simulation in a volume averaged frame where *α* is similar to the SUSHI frame (measured *α* = 0.27) but there is no structure (**Figure 4A**).

**Figure 4.**
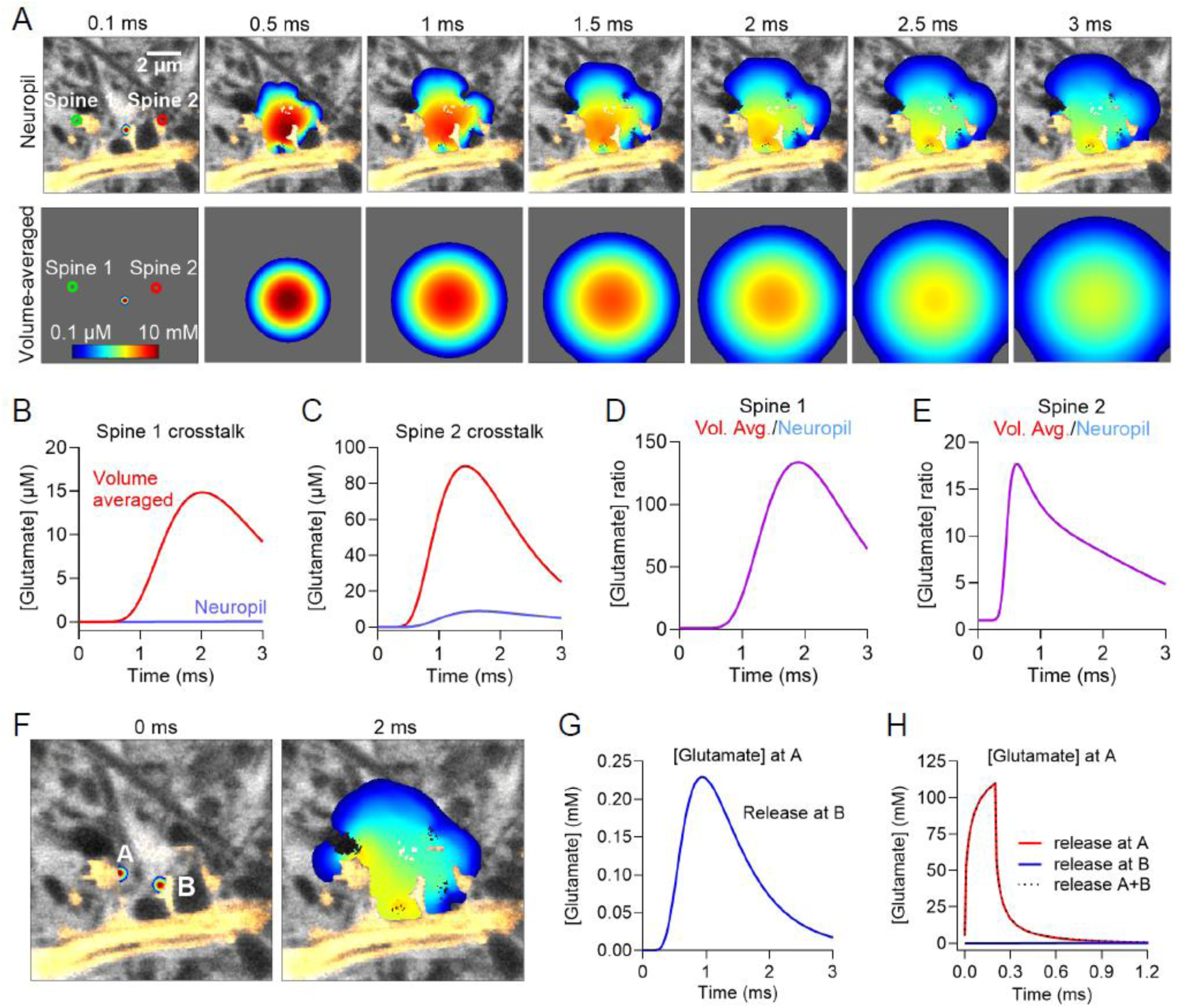
Dendritic spine geometry facilitates extrasynaptic diffusional clearance (**A**) Simulation of synaptic glutamate release at a dendritic spine with two immediate neighboring spines 1 and 2, and the corresponding frame volume-averaged to eliminate any structural information. (**B**) Peak glutamate reaching neighboring Spine 1 through synaptic crosstalk is 14.8 μM in the case of volume averaging, while in the SUSHI image with neuropil context the maximum reached concentration is merely around 0.04 μM. (**C**) Similarly for neighboring Spine 2, in the SUSHI image case the maximum crosstalk reaches 8.8 μM while in the volume-averaged case it is 89.5 μM. The ratios of crosstalk in the volume averaged and neuropil context amounted to 134 for Spine 1(**D**) and 18.0 for Spine 2(**E**), suggesting that in these two cases the neuropil greatly reduces crosstalk compared to the volume averaged scenario.(**F**) Moving the neighbor Spine A synapse even closer to the source spine B, we analyzed simulated glutamate concentrations at A. (**G**) Crosstalk now reached 0.22 mM. (**H**) Still this crosstalk appeared negligible compared to the concentrations reached after direct glutamate signaling onto A where concentrations surpassed 100 mM.

We observed that having knowledge on the ECS geometric context would dramatically affect the resulting crosstalk, and that diffusional clearance appeared faster in the neuropil context than predicted from volume-averaged data. Notably, neighboring Spine 1 would see more than 100-fold higher glutamate concentrations in the volume averaged case, while neighboring Spine 2 would see more than 10-fold higher concentrations in the volume averaged case (**Figure 4B-E**). This suggests that for the given case the perisynaptic spine ECS geometry facilitates synaptic clearance and minimizes extracellular crosstalk beyond the levels predicted based on averaged data. We wondered how this level of crosstalk would compare to the glutamate concentration a synapse sees upon direct signalling, and to our surprise the added effect of crosstalk for a 1 μm distanced spine pair turned out to be practically indiscernible from direct signalling alone, suggesting the crosstalk effect is in this case and context negligible (**Figure 4F-H**). While this is simply one example, it examplifies how glutamatergic synaptic crosstalk can be shaped by ECS geometry to be more than 100-fold less than predicted from volume averaged data, which would reduce signaling noise and enhance fast independent signalling at glutamatergic synapses.

### ECS geometry facilitates extrasynaptic tonic inhibition from diffusional GABA synapse spillover

We turned from protruding glutamatergic spine synapses to membrane-level somatodendritic GABAergic synapses to learn how ECS geometry here influences perisynaptic diffusion. Vesicular GABA release was simulated in a SUSHI image at 47 distributed somatodendritic and dense neuropil release points separated by around 5 μm from each other and using *DGABA* = 1.1 x 10^-9^ m^2^/s (Rodrigo et al., 2017; Khanal et al., 2022) (**Figure 5A**). The chosen synapse density is in the range of around 0.1 per µm repoted for CA1 pyramidal cells(Tasciotti et al., 2025). At each site 10,000 molecules were released at 10, 25, 50 and 100 Hz that is within the published firing ranges of GABAergic interneurons (He et al., 2021). Again, the results were compared to a corresponding homogenized image that has the same average α but discards ECS structure (**Figure 5B**).

**Figure 5.**
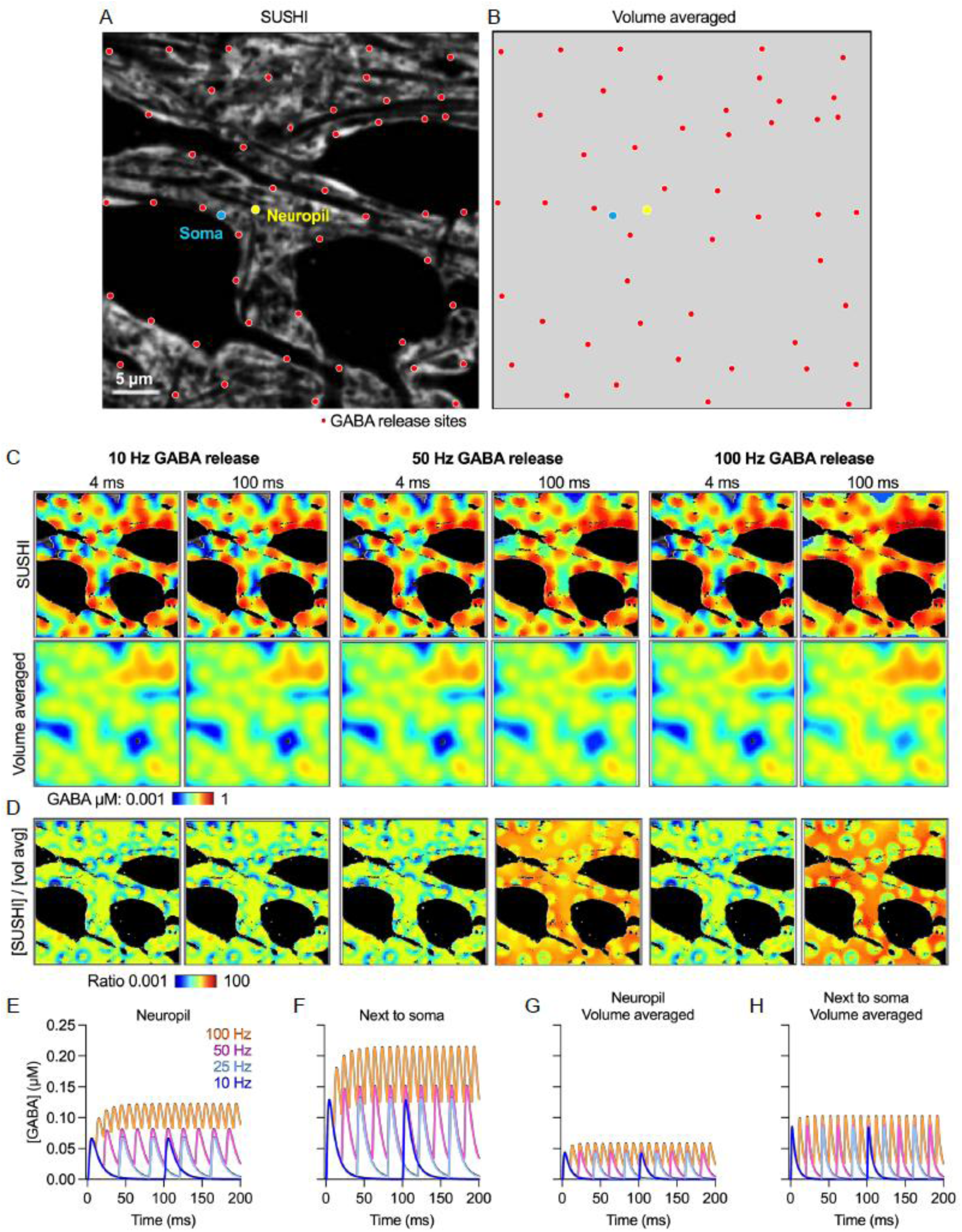
ECS geometry shapes GABA spillover to enhance extrasynaptic inhibition. (**A, B**) SUSHI image and the equivalent volume-averaged image. The images show GABA release sites distribute 5 μm away from each other (red dots), a measuring point next to the soma (blue) and in the neuropil (yellow). (**C**) Representative simulations of GABA release at 10 Hz, 25Hz, 50 Hz and 100 Hz. Images show the GABA distribution after 4 and 100 ms for the SUSHI and the volume-averaged image. (**D**) Ratio between the observed concentrations in the SUSHI and volume-averaged image. The ratio is displayed for 10Hz, 50 Hz and 100 Hz GABA release frequencies, at 4ms and 100ms into the stimulation sequence. (**E, F**) Extracellular GABA concentration measured over time in neuropil and next to the soma in the SUSHI image, and correspondingly in the volume-average image (**G, H**).

We observed in the SUSHI image simulations that at increasing release frequencies the extracellular GABA concentration plateau grows to reach a higher steady-state (**Figure 5C- F**). This increase was more pronounced along somatic cell membranes of the receiving neuron, indicating that the large smooth ECS geometry at the soma membrane surface facilitates lateral diffusion from somatodendritic synapses to nearby extrasynaptic receptors, corroborating our tensor mapping above (**Figure 5E-F**). Measuring the corresponding concentrations at the same sites in the volume averaged image without ECS structure, this build-up of extracellular GABA was manifold less pronounced, highlighting the applicability of *DifFlux* for nano- and microscale simulations (**Figure 5G-H**).

Our results suggest that somatodendritic synapse geometry facilitates lateral diffusion along the membrane and thus spillover to perisynaptic receptors and extrasynaptic GABAergic inhibition. Interestingly, it suggests that for regular release frequencies steady state GABA concentration gradients can build up and extend from the soma surface and into the neuropil, meaning that extrasynaptic receptors on the cell membrane see a higher concentration of GABA than predicted from considering average interstitial GABA concentrations. Again, the steepness and stability of such steady state gradients will be hugely context dependent.

## Discussion

There is still no straightforward approach to empirically measure point-source diffusion in spatially resolved brain ECS geometries and working with diffusion from the stance of volume-averaged data is currently the norm, as recently reviewed (Soria et al., 2020). Existing mathematical and computational diffusion models are accordingly confounded by relying on synthetic ECS lattice geometries that are designed to reproduce volume-averaged empirical data (Syková and Nicholson, 2008). One exception utilized binarized and volume-rendered fixed tissue geometries generated from 3D electron microscopy images, though anisotropy or anomalous diffusion was not analyzed. Also, the necessity for offline tissue volume corrections in this approach may introduce structural ECS artifacts (Kinney et al., 2013). The highest resolution empirical approach is single molecule tracking of diffusing single-walled carbon nanotubes that yield both geometric and diffusional data, though a key limitation is that single carbon nanotubes do not distribute homogenously and exhaustively in brain parenchyma, which limits what can be learned about the relationship between ECS geometry and diffusion, particularly about local point-source diffusion (Godin et al., 2017). Another notable technique is 2-photon microscopy based time-resolved fluorescence anisotropy imaging that also yields simultaneous diffusional and geometric data, though it lacks the resolution to resolved the brain neuropil, and is restricted in the ranges of diffusion it can resolve (Zheng et al., 2017). It would be interesting to apply this approach based on STED microscopy to effectively reconcile nanoscale resolution imaging with concurrent ECS viscosity data.

Our computational *DifFlux* and Monte Carlo models based on perfusion labelling and super-resolved fluorescence microscopy of the ECS in brain slices allows nanoscale analysis of molecular diffusion in actual images of live brain parenchyma. Instead of an image binarization approach and the associated confounders of thresholding and 3D reconstructions, it adopts a largely analogue approach in raw fluorescence images and exploits the full pixel bit depth to harness 3D information about ECS/cellular structure ratios in individual pixels. While higher spatial resolution gives more accurate results, the model does not necessitate resolving ECS geometry as long as a qualified estimate or measurement of background and maximum observable ECS fluorescence can be made. The model thus requires few underlying assumptions, making it robust and applicable across any fluorescence microscopy modality where background and maximum fluorescence input parameters can be provided. It simulates diffusion in 3D and can be applied in 3D images, as well as 2D images where binarized models fall short because they produce closed maze geometries.

Our modeling approach is purely structural and blind to ECS viscosity effects or binding/uptake of the diffusing molecular species, though both parameters can conceivably be incorporated if experimenters have knowledge about these. Indeed, this could be done in the same SUSHI images by retrospectively mapping onto these immunohistochemically mapped matrix constituents, membrane receptors, transporters, or other targets of interest.

We acknowledge that there are further confounders of our model. Firstly, the background fluorescence across images is expectedly not fully homogenous, leading to an error in our estimates of *α* based on *FPix* that will scale with our signal to noise ratio. This error is expectedly low because of the high contrast of the general volume labelling approach, and introducing heterogeneous background correction complicates the modeling beyond the current study. Further, the microscope imaging PSF is not entirely homogenous, and a given pixel intensity may therefore not scale completely linearly with pixel *α*. Though we are blind to which part of the PSF is actually sampling ECS vs cellular space, and the resulting error will expectedly be symmetric and not biased toward lower or higher pixel values. The same confounder applies to our image border escape coefficients in the x,y,z planes, where this is not in reality homogeneous even if we assume it in the model. This is especially true for the z-axis where we for 3Dsmpl based analyses use a predetermined escape factor to simulate escape out of a given x,y plane, though our analyses suggest that results are comparable to true 3D modelling, which likely reflects that the contribution of molecular “bounce” between z-planes is low compared to the contribution of molecules moving within a given x,y plane. Indeed, our relatively modest z-resolution here adds robustness to the model, as much more molecules are transitioning between neighboring pixels in the x,y plane than in and out of z-planes for a given model time step. These confounders presumably add noise without biasing our results, and we accept them *as are* to have an as simple as possible model with minimal assumptions. Running the *DifFlux* model in parallel to Monte-Carlo simulations applied in a corresponding analogue manner yielded comparable results. Whereas *DifFlux* represents a bulk model that does not consider trajectories of individual molecules, the Monte Carlo approach allows tracking of individual molecules and analysis of MSD. The advantage of *DifFlux* here is that it is much lighter computationally, though the two are highly complementary.

Having built a diffusion model and gained confidence in its performance to reveal diffusion gradients in SUSHI images of live brain tissue ECS, we applied *DifFlux* to simulate point-source release of transmitters in the neuropil. We observed anomalous sub-diffusion compared to free diffusion, as expected and observed by others (Grassi et al., 2023). We went on to find pronounced anisotropic diffusion on the micron scale, which would cancel out on larger scales that are within the reach of volume-averaging techniques. These findings draw into question the soundness of applying volume-averaged diffusion models to understand ECS diffusion on single-micron scales where volume-transmitters conceivably will exert their main effect. We confined our modeling to relatively homogeneous CA1 *stratum radiatum* ECS, and we expect that modelling diffusion across cell layers of brain regions would identify anisotropic diffusion also at larger scales that encompass distinct cell layers, vessels, or ventricular structures.

We further analyzed directionality of diffusion tensors with reference to ECS geometries and observed that dendrites facilitate diffusion along their direction, while dense neuropil imposes no apparent diffusional directionality. Here, directionality results will depend on the exact location of the point-sources observed and the running time of individual simulations. We note that this is not merely a technical aspect as it will also apply to biological processes, where the degree of diffusion anomaly, anisotropy and directionality will depend on the diffusion distance and time from a given point-source event.

The observation of differences in diffusion along larger membranes and in denser neuropil prompted us to simulate perisynaptic diffusion of transmitters around respective dendritic spines that effectively position the synapse in dense neuropil, and somatodendritic synapses that are flush with somatic or dendritic membranes.

Our modelling found that transmitter clearance from glutamatergic synapses on dendritic spines is facilitated by spine geometry to minimize extracellular synaptic crosstalk and that clearance is higher than predicted from volume averaged models. This suggests that dendritic spine geometry contribute in diffusional clearance from the synaptic and perisynaptic ECS and supports the common view of glutamatergic synapses as individual, independent, fast-tunable substrates of memory and learning processes (Sala and Segal, 2014; Tønnesen and Nägerl, 2016). Conversely, we found that the somatodendritic location of GABAergic synapses facilitate lateral diffusion to nearby extra-synaptic receptors on the same membrane, meaning they will locally augment tonic GABAergic inhibition emerging from a given combination of released GABA molecules and extra-synaptic receptors. Interestingly, this direct signaling regulation can happen independently of presynaptic quantal size, GABA release frequency, transporter activity, post-synaptic receptor numbers and states, which are the variables by which we currently understand and investigate extrasynaptic signal regulation. In other words, this regulation mechanism that can operate independently of the transmitting and receiving cell pair, and will expectedly be influential in terms of neuronal and circuit excitability. This suggests GABA synapse geometry has evolved to facilitate synaptic as well as spill-over derived extrasynaptic signaling.

Our data add a new ECS perspective to previous seminal work concluding that tonic inhibition of hippocampal granule cells emerges primarily from somatic post-synapses because it persists when trimming away the dendritic tree (Soltesz et al., 1995). Our results imply that even for homogenously distributed synapses across granule dendrites and soma, tonic inhibition of the somatic compartment is higher based on the very dense granule cell layer having a low ECS volume fraction and diffusion being highly restricted among the dense soma layer which would result in higher GABA concentrations than predicted for the molecular layer (Tønnesen et al., 2018).

In concert, our perisynaptic diffusion modelling suggests the very different glutamatergic and GABAergic synapse geometries support separate functions and increase metabolic and computational efficiency in a transmitter system specific manner. This should be thought of in terms of probabilities, rather than as a ubiquitous truth, and our mission here is to point to new potential functions of the ECS geometry that are not uncovered by lower resolution volume-averaging approaches.

Down the line, our model will allow us and others to further understand how brain ECS diffusional properties may change across brain regions, physiological conditions such as development and ageing, and how inflammation or neurodegenerative disorders associated with extracellular protein aggregation impact not only signaling, but also metabolite clearance *via* the ECS.

We already model diffusion from repeated release over time, though it will be interesting to extend the *DifFlux* model to accept 4D SUSHI images and look at ECS structural dynamics, and additionally combine the 4D modelling predictions with functional data, such as calcium-imaging or electrophysiology experiments to further explore how ECS geometry may impact brain function across scales. And by adding data from immunohistochemistry, fluorescence correlation spectroscopy, polarization microscopy, and more, we can likely improve the model by incorporating non-structural regulators of ECS diffusion, for example viscosity and molecular binding data. The *DifFlux* model could also be applied to simulate diffusion in other tissue than brain, and to simulate intracellular diffusion rather than that through the ECS.

## Acknowledgements

JT and PG acknowledge funding support from the Aligning Science Across Parkinson’s initiative (ASAP-020505), from the Spanish Government (PID2020-113894RB-I00, CNS2022-135903), including funding through the Spain-US Collaborative Research in Computational Neuroscience (CRCNS) program (PCI2022-135040-2) and resources from Spain’s National Recovery and Resilience plan financed by the Next Generation EU (NGEU) instrument. PG was initially hired through a University of the Basque Country UPV/EHU doctoral program grant (GIU21/048). We further acknowledge support from The Sigma Xi Grants in Aid of Research (GIAR) program. FS and RS supported by NIH-NINDS R01NS130759, NSF-EFRI BEGIN OI 2515404, NSF-EFRI BRAID 2318139.

## Data availability

All simulations were programmed in MATLAB (Natick, MA). The code and analyses are available at <u>Github.com/TonnesenLab/Diffusion-Model</u> and <u>Github.com/Santamarialab.</u>

All images and results are available at the Zenodo repository (DOI: 10.5281/zenodo.17473125).

## Key Resources Table

| Reagents | Provider | Identifier |
| --- | --- | --- |
| Hank's balanced salt solution (HBSS) | Thermo-Fisher Scientific | 24020091 |
| Minimum Essential Media (MEM) | Thermo-Fisher Scientific | 11095080 |
| L-Glutamine | Thermo-Fisher Scientific | 25030024 |
| B27 supplement | Thermo-Fisher Scientific | 17504-044 |
| Penicillin-Streptomycin | Thermo-Fisher Scientific | 15140122 |
| Horse serum | Thermo-Fisher Scientific | 26050088 |
| Glucose | Sigma-Aldrich | G7021 |
| Kynurenic Acid | Sigma-Aldrich | K3375 |
| HEPES | Sigma-Aldrich | H3375-100G |
| poly-L-lysine | Sigma-Aldrich | P8920 |
| Chicken Plasma | Sigma-Aldrich | P3266 |
| Sucrose | Sigma-Aldrich | S0389 |
| Thrombin | Sigma-Aldrich | 112374 |
| Ascorbic L-acid | Sigma-Aldrich | A4403 |
| Uridine | Sigma-Aldrich | U3750 |
| Ara-C | Sigma-Aldrich | C6645 |
| 5-Fluoro-2'-deoxyuridine | Sigma-Aldrich | F0503 |
| NaCl | Sigma-Aldrich | S7653 |
| KCl | Sigma-Aldrich | P5405 |
| CaCl<sub>2</sub> | Sigma-Aldrich | C8106 |
| MgCl<sub>2</sub> | Sigma-Aldrich | M8266 |
| Trolox | Sigma-Aldrich | 238813 |
| Calcein disodium salt | Sigma-Aldrich | 21030 |
| D-Mannitol | Sigma-Aldrich | M4125 |
| Software | Provider | Identifier |
| Matlab R2019b | MathWorks | www.mathworks.com |
| Prism 10 | GraphPad | www.graphpad.com |
| Image J 1.53e | ImageJ | https://imagej.net |

**Supplementary Figure 1.**
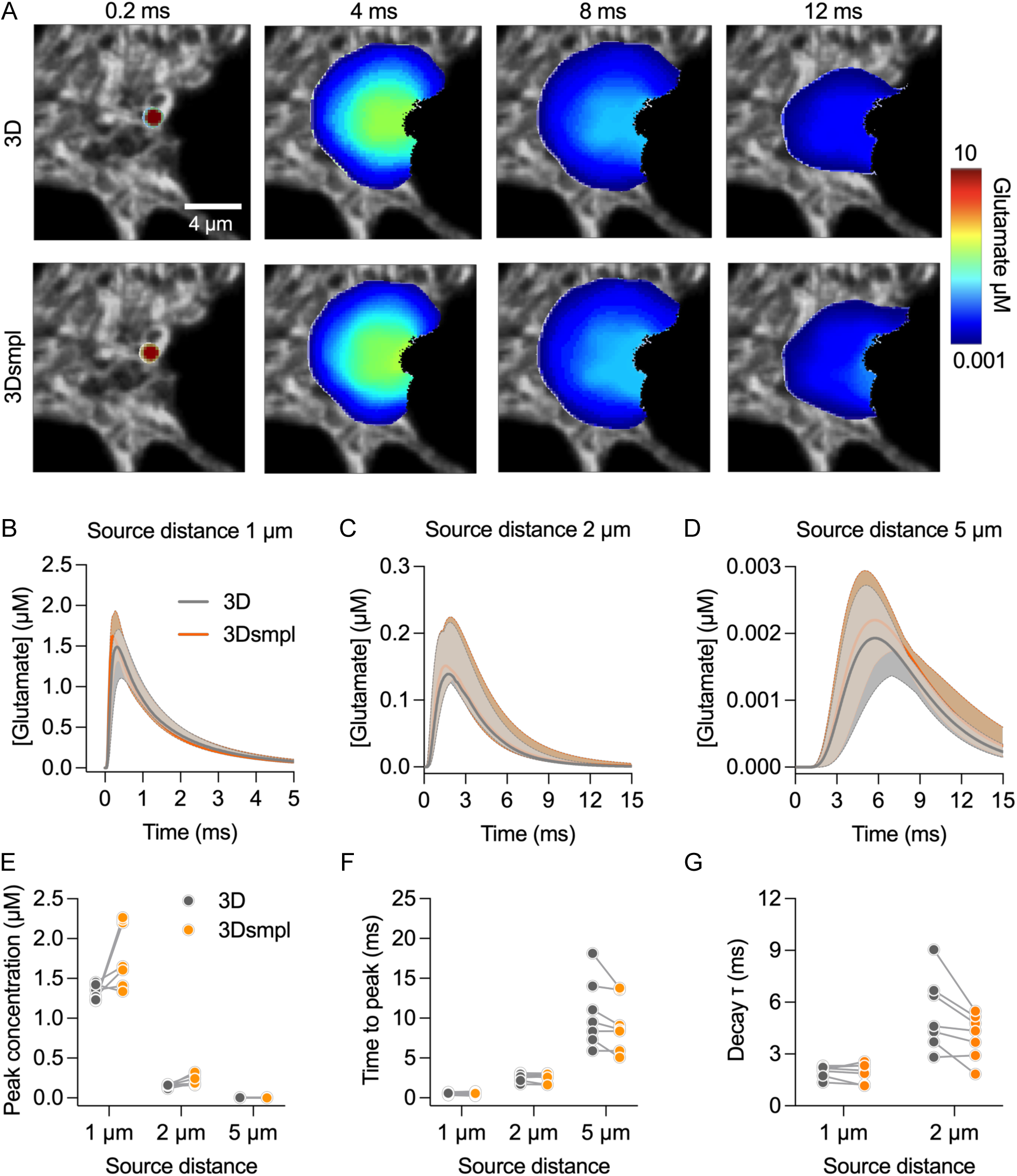

**Supplementary Figure 2.**
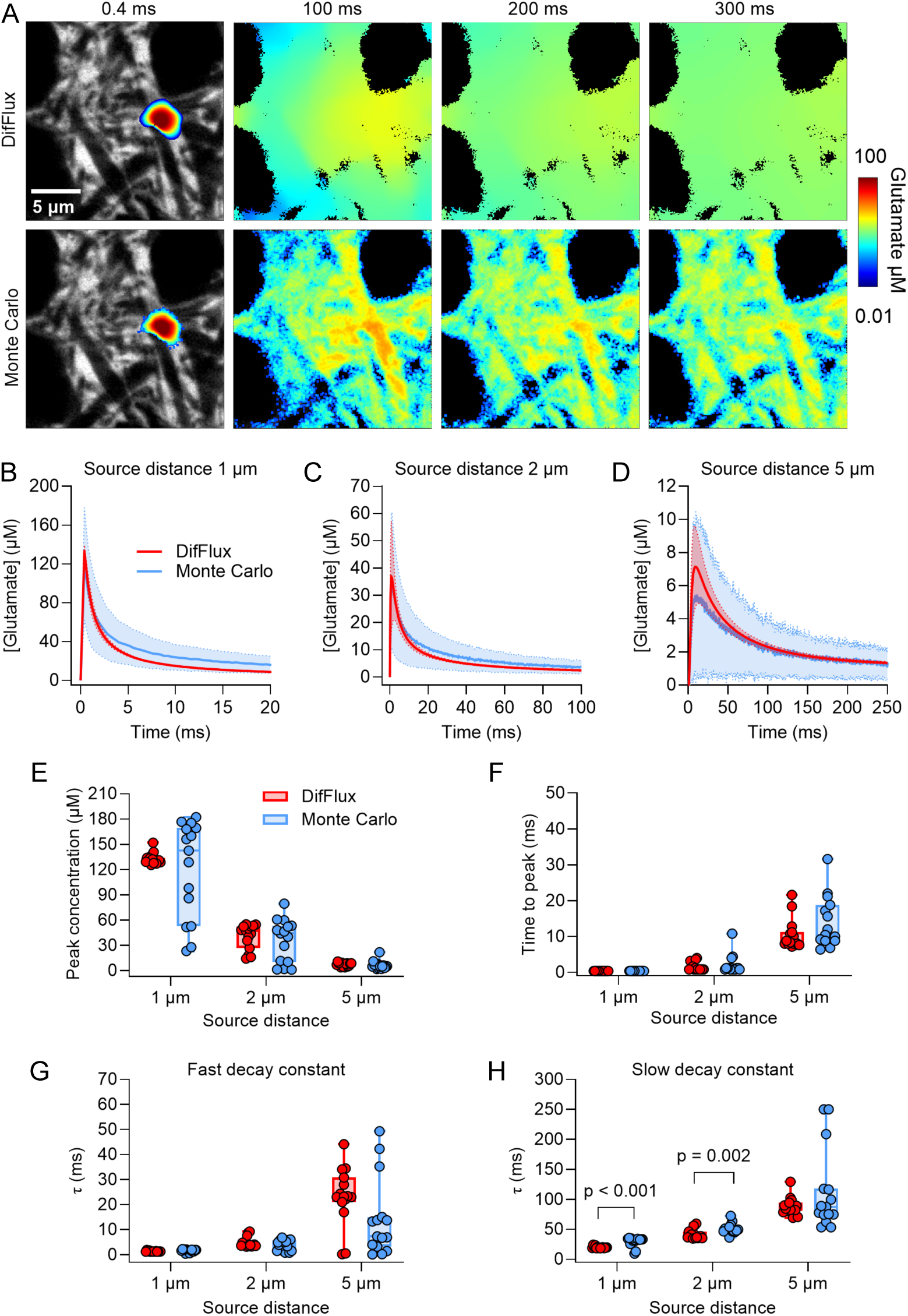

## Notes

https://zenodo.org/records/17473125

### Competing Interest Statement

The authors have declared no competing interest.

