## Supplementary Material for "Extracellular space diffusion modelling identifies distinct functional advantages of archetypical glutamatergic and GABAergic synapse geometries"

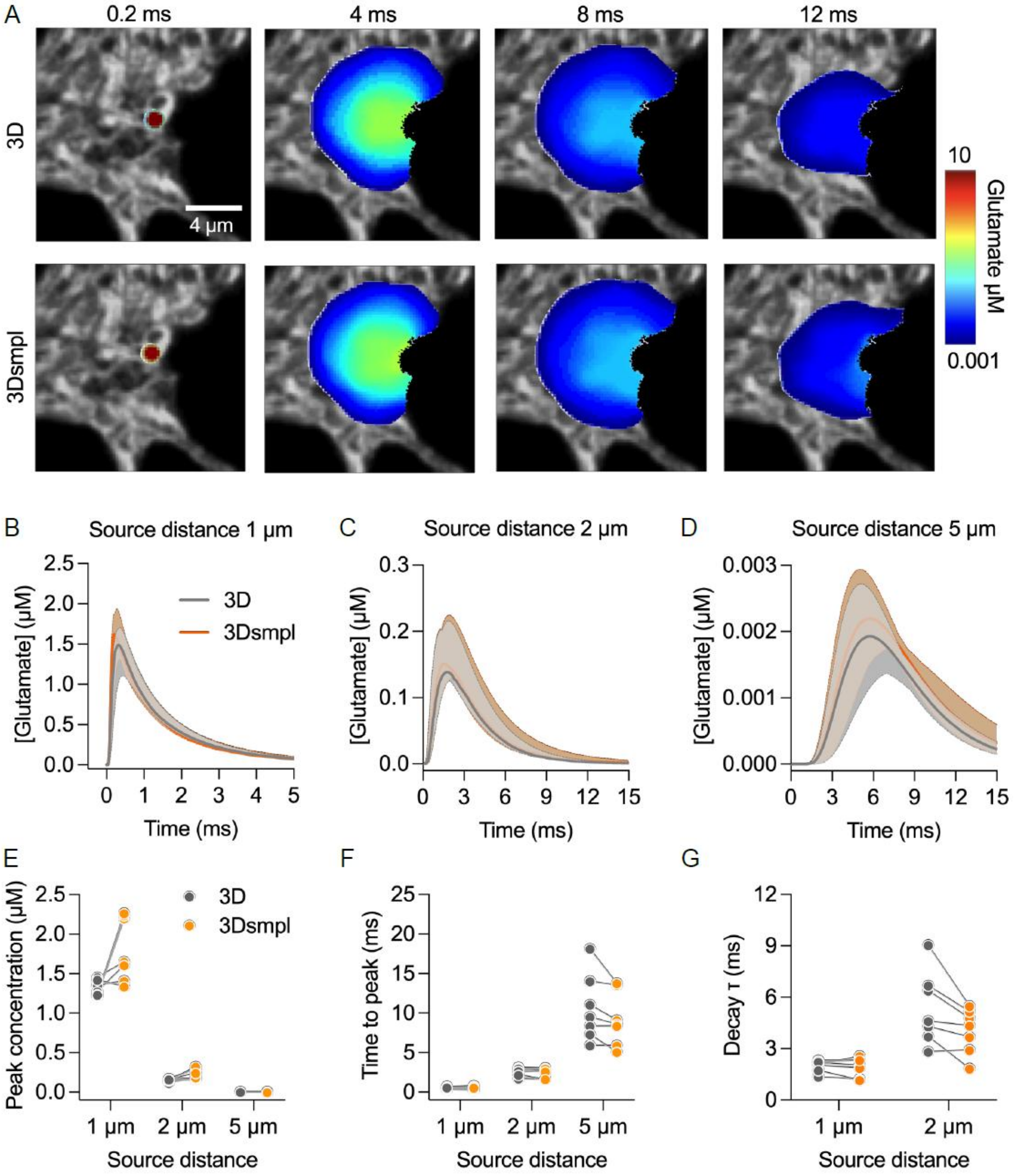


***Suppl. Figure 1. Validating the simplified 3D$_{smpl}$ diffusion model***
*(**A**) Comparison example of 3D and 3D$_{smpl}$ glutamate diffusion modelling observed in a single xy plane based on real 3D tissue geometry in a SUSHI z-stack and applying a constant z-axis escape factor (3D$_{smpl}$) to the xy plane and disregarding the other planes of the stack. (**B-D**) Resulting glutamate concentrations measured at fifteen equally spaced concentric locations at 1 µm, 2 µm and 5 µm from the point-source. Graphs display mean with SD (n = 15 measuring points). (**E-G**) Peak concentration, time to peak and decay constant (τ). Graphs display the mean values for the 3D simulations in each of the SUSHI z-stack and their corresponding 3D$_{smpl}$ simulation in the middle plane. (n = 7 z-stacks). There was no significant difference in the observed peak concentrations, (G) nor in the time to reach the peak concentration, or decay constant between the two approaches. Numerical values are listed in Supplementary Tables 2, 3 and 4.*

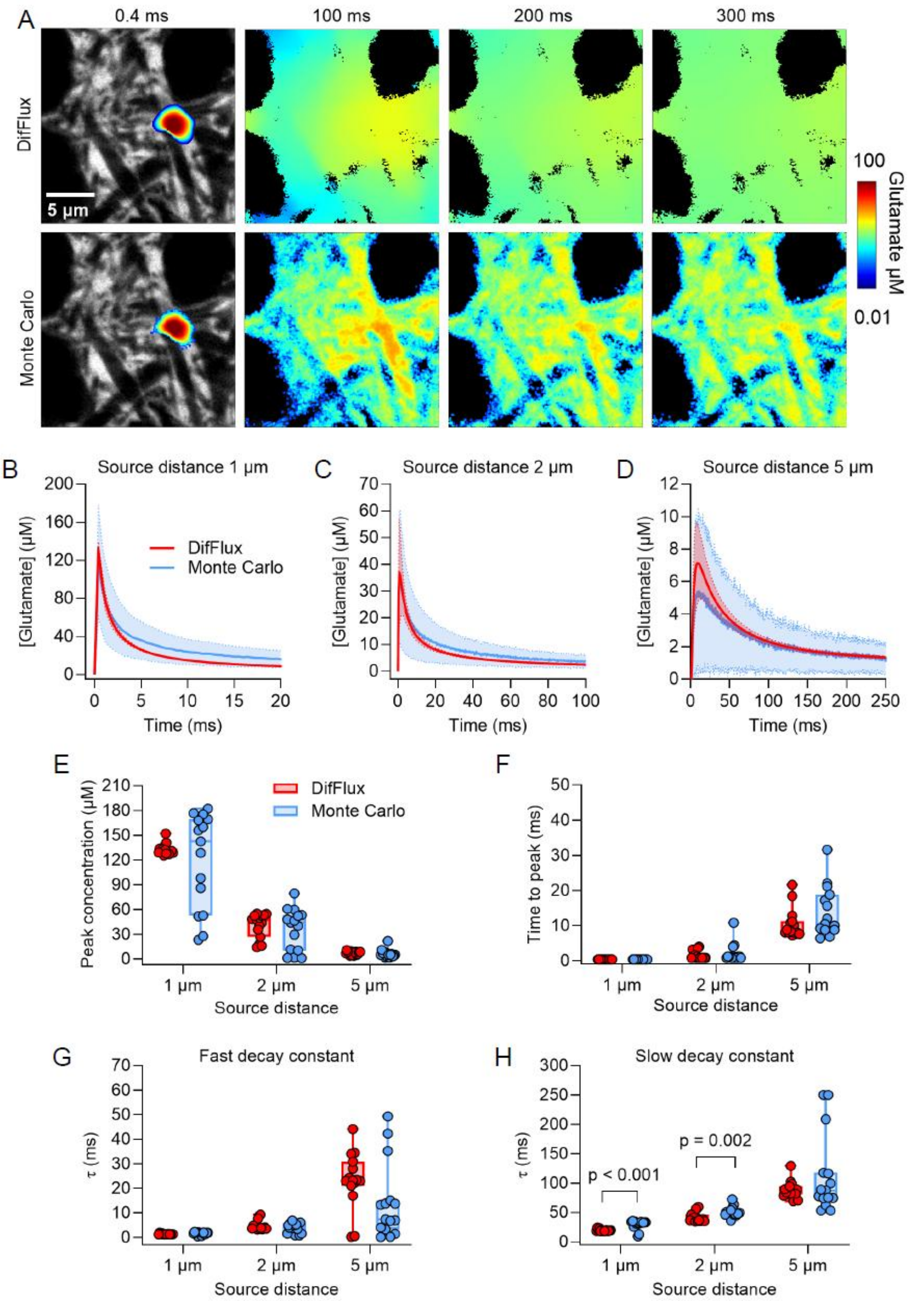


***Suppl. Figure 2. DifFlux and Monte Carlo simulations yield comparable ECS diffusion results***

*(**A**) Modelling point-source release of 100.000 particles by DifFlux and Monte Carlo simulations in the same SUSHI frame of neuropil. (**B**) Resulting concentrations profiles over time at 1 µm, 2 µm and 5 µm source distance measured at fifteen equally spaced concentric locations for each distance. Graphs depict mean with SD. There were no significant differences in peak concentrations at 1 µm, 2 µm and 5 µm from the source (graphs show mean with SD) (**E**), nor in corresponding concentration peak times measured (Bars show range, median and quartiles) (**F**). Fitting a double-exponential function to the decay phase of the diffusion gradient revealed no differences in the fast component (**G**), while at 1 µm and 2 µm from the source there were significant distances between the DifFlux and Monte Carlo simulations source (Multiple Mann-Whitney test), even if the numerical difference was modest (**H**). See Supplementary Tables 5 and 6 for numerical results.*

**Suppl. Table 1. Radially Averaged power spectral density (PSD) and mean square displacement ($<r^2>$) from Monte Carlo simulated diffusion**

| | Stack 1 | Stack 2 | Stack 3 | Stack 4 | Stack 5 | Stack 6 | Stack 7 | Overall |
|---|---|---|---|---|---|---|---|---|
| PSD β (low frequencies) | 1.85 $R^2 = 0.99$ | 3.64 $R^2 = 0.96$ | 1.97 $R^2 = 0.98$ | 1.87 $R^2 = 0.96$ | 2.07 $R^2 = 0.95$ | 2.39 $R^2 = 0.98$ | 1.87 $R^2 = 0.97$ | 2.04 $R^2 = 0.99$ |
| PSD β (high frequencies) | 4.07 $R^2 = 0.99$ | 3.70 $R^2 = 0.99$ | 3.80 $R^2 = 0.99$ | 3.66 $R^2 = 0.99$ | 3.06 $R^2 = 0.99$ | 3.25 $R^2 = 0.98$ | 3.49 $R^2 = 0.98$ | 3.62 $R^2 = 0.99$ |
| $< r^2 >$ γ mean ± SD | 0.96 ± 0.03 | 0.81 ± 0.04 | 0.86 ± 0.07 | 0.92 ± 0.03 | 0.87 ± 0.05 | 0.81 ± 0.06 | 0.95 ± 0.04 | 0.88 ± 0.05** |

β denotes PSD fit exponent, γ denotes $< r^2 >$ fit exponent.

** Statistical difference between mean γ and 1, p<0.0001 (t-test, Wilcoxon).

**Suppl. Table 2. Comparing 3D and 3D*smpl* *DifFlux* approaches resulting peak concentrations (Mean ± SD, n = 15).**

| **Peak concentration (µM) at 1 µm from source** | | | | | | | |
|---|---|---|---|---|---|---|---|
| | Stack 1 | Stack 2 | Stack 3 | Stack 4 | Stack 5 | Stack 6 | Stack 7 |
| 3D | 1.36 ± 0.27 | 1.45 ± 0.19 | 1.42 ± 0.16 | 1.27 ± 0.37 | 1.22 ± 0.34 | 1.24 ± 0.20 | 1.23 ± 0.19 |
| $3D_{smpl}$ | 1.41 ± 0.27 | 1.65 ± 0.21 | 1.34 ± 0.17 | 1.61 ± 0.18 | 2.20 ± 0.44 | 2.24 ± 0.33 | 2.26± 0.27 |
| **Peak concentration (µM) at 2 µm** | | | | | | | |
| | Stack 1 | Stack 2 | Stack 3 | Stack 4 | Stack 5 | Stack 6 | Stack 7 |
| 3D | 0.14 ± 0.05 | 0.15 ± 0.03 | 0.17 ± 0.03 | 0.17 ± 0.06 | 0.12 ± 0.02 | 0.15 ± 0.03 | 0.16 ± 0.03 |
| $3D_{smpl}$ | 0.17 ± 0.06 | 0.16 ± 0.3 | 0.18 ± 0.03 | 0.30 ± 0.08 | 0.19 ± 0.02 | 0.32 ± 0.06 | 0.24 ± 0.04 |
| **Peak concentration (µM) at 5 µm** | | | | | | | |
| | Stack 1 | Stack 2 | Stack 3 | Stack 4 | Stack 5 | Stack 6 | Stack 7 |
| 3D | 0.005 ± 0.002 | 0.002 ± 0.0004 | 0.004 ± 0.002 | 0.006 ± 0.002 | 0.001 ± 0.001 | 0.002 ± 0.001 | 0.003 ± 0.0004 |
| $3D_{smpl}$ | 0.003 ± 0.001 | 0.002 ± 0.0004 | 0.002 ± 0.001 | 0.006 ± 0.002 | 0.001 ± 0.0004 | 0.005 ± 0.001 | 0.001 ± 0.0002 |

**Suppl. Table 3. Comparing 3D and 3D*smpl* *DifFlux* approaches resulting time to peak concentration (Mean ± SD, n = 15).**

| **Time to peak (ms) at 1 µm from source** | | | | | | | |
|---|---|---|---|---|---|---|---|
| | Stack 1 | Stack 2 | Stack 3 | Stack 4 | Stack 5 | Stack 6 | Stack 7 |
| 3D | 0.52 ± 0.16 | 0.34 ± 0.05 | 0.41 ± 0.05 | 0.56 ± 0.24 | 0.73 ± 0.15 | 0.64 ± 0.13 | 0.58 ± 0.11 |
| $3D_{smpl}$ | 0.56 ± 0.18 | 0.36 ± 0.04 | 0.40 ± 0.06 | 0.68 ± 0.29 | 0.79 ± 0.16 | 0.70 ± 0.16 | 0.56± 0.10 |
| **Time to peak (ms) at 2 µm** | | | | | | | |
| | Stack 1 | Stack 2 | Stack 3 | Stack 4 | Stack 5 | Stack 6 | Stack 7 |
| 3D | 3.12 ± 1.41 | 1.72 ± 0.20 | 2.88 ± 0.87 | 2.56 ± 0.88 | 2.97 ± 0.57 | 2.65 ± 0.63 | 2.19 ± 0.32 |
| $3D_{smpl}$ | 3.056 ± 1.59 | 1.66 ± 0.25 | 2.73 ± 0.92 | 2.65 ± 0.76 | 2.97 ± 0.71 | 2.61 ± 0.71 | 1.63 ± 0.21 |
| **Time to peak (ms) at 5 µm** | | | | | | | |
| | Stack 1 | Stack 2 | Stack 3 | Stack 4 | Stack 5 | Stack 6 | Stack 7 |
| 3D | 11.05 ± 2.63 | 5.89 ± 0.68 | 14.01 ± 3.97 | 18.12 ± 2.11 | 9.49 ± 1.78 | 8.35 ± 1.83 | 7.31 ± 0.50 |
| $3D_{smpl}$ | 9.11 ± 2.72 | 5.90 ± 0.88 | 13.56 ± 8.32 | 13.79 ± 2.57 | 8.61 ± 2.48 | 8.38 ± 2.64 | 5.08 ± 0.63 |

**Suppl. Table 4. Comparing 3D and 3D$_{smpl}$ *DifFlux* approaches resulting decay constants (Mean ± SD, n = 15).**

| τ (ms) at 1 μm from source | | | | | | | |
|---|---|---|---|---|---|---|---|
| | Stack 1 | Stack 2 | Stack 3 | Stack 4 | Stack 5 | Stack 6 | Stack 7 |
| 3D | 2.32 ± 0.36 | 1.35 ± 0.06 | 2.18 ± 0.21 | 2.06 ± 0.47 | 2.01 ± 0.25 | 2.23 ± 0.20 | 1.74 ± 0.21 |
| 3Dsmpl | 2.31 ± 0.34 | 1.23 ± 0.09 | 2.04 ± 0.14 | 2.56 ± 0.30 | 1.88 ± 0.21 | 2.33 ± 0.15 | 1.16 ± 0.07 |
| **τ (ms) at 2 μm** | | | | | | | |
| | Stack 1 | Stack 2 | Stack 3 | Stack 4 | Stack 5 | Stack 6 | Stack 7 |
| 3D | 6.39 ± 1.17 | 2.82 ± 0.20 | 6.68 ± 1.35 | 9.04 ± 2.00 | 4.29 ± 0.96 | 4.61 ± 0.78 | 3.70 ± 0.38 |
| 3Dsmpl | 4.78 ± 1.30 | 2.91 ± 0.44 | 5.17 ± 1.37 | 5.48 ± 0.70 | 3.68 ± 1.02 | 4.34 ± 0.87 | 1.83 ± 0.14 |

**Suppl. Table 5. Comparing *DifFlux* and Monte Carlo simulations resulting peak concentrations (Mean ± SD, n = 15)**

| Source distance (μm) | *DifFlux* peak concentration (μM) | MC peak concentration (μM) | *DifFlux* time to peak (ms) | MC time to peak (ms) |
|---|---|---|---|---|
| 1 | 132.9 ± 6.6 | 119.9 ± 58.0 | 0.4 ± 0 | 0.4 ± 0 |
| 2 | 40.5 ± 14.8 | 36.0 ± 25.1 | 1.47 ± 1.14 | 2.08 ± 2.67 |
| 5 | 7.3 ± 2.2 | 6.4 ± 5.0 | 10.43 ± 4.24 | 13.89 ± 7.07 |

**Suppl. Table 6. Comparing *DifFlux* and Monte Carlo simulations resulting decay constants (Mean ± SD, n = 15)**

| Source distance (μm) | *DifFlux* fast decay constant (ms) | MC fast decay constant (ms) | *DifFlux* slow decay constant (ms) | MC slow decay constant (ms) |
|---|---|---|---|---|
| 1 | 1.37 ± 0.21 | 1.59 ± 0.55 | 19.84 ± 1.73 | 29.66 ± 7.89** |
| 2 | 4.83 ± 2.02 | 3.80 ± 2.07 | 42.04 ± 8.57 | 51.21 ± 8.33* |
| 5 | 23.29 ± 11.50 | 14.19 ± 15.59 | 88.17 ± 14.58 | 112.61 ± 67.33 |

** Statistical difference between *DifFlux* and MC $p < 0.001$

* Statistical difference between *DifFlux* and MC $p = 0.002$
